# Fine-grained topographic organization within somatosensory cortex during resting-state and emotional face-matching task and its association with ASD traits

**DOI:** 10.1101/2022.04.26.489525

**Authors:** Christina Isakoglou, Koen V. Haak, Thomas Wolfers, Dorothea L. Floris, Alberto Llera, Marianne Oldehinkel, Natalie J. Forde, Bethany F. M. Oakley, Julian Tillmann, Rosemary J. Holt, Carolin Moessnang, Eva Loth, Thomas Bourgeron, Simon Baron-Cohen, Tony Charman, Tobias Banaschewski, Declan G. M. Murphy, Jan K. Buitelaar, Andre F. Marquand, Christian F. Beckmann, the EU-AIMS LEAP Group

## Abstract

**BACKGROUND:** Sensory atypicalities are particularly common in autism spectrum disorders (ASD). Nevertheless, our knowledge about the divergence of the underlying somatosensory region and its association with ASD phenotype features is limited.

**METHODS:** We applied a data-driven approach to map the fine-grained variations in functional connectivity of the primary somatosensory cortex (S1) to the rest of the brain in 240 autistic and 164 neurotypical individuals from the EU-AIMS LEAP dataset, aged between 7 and 30. We estimated the S1 connection topography (‘connectopy’) during rest and during the emotional face-matching (Hariri) task, an established measure of emotion reactivity, and accessed its association with a set of clinical and behavioral variables.

**RESULTS:** We demonstrated that the S1 connectopy is organized along a dorsoventral axis, mapping onto the somatotopic organization of S1. We found that its spatial characteristics were linked to the individuals’ adaptive functioning skills, as measured by the Vineland Adaptive Behavior Scales, across the whole sample. Higher functional differentiation characterized the S1 connectopies of individuals with higher daily life adaptive skills. Notably, we detected significant differences between rest and the Hariri task in the S1 connectopies, as well as their projection maps onto the rest of the brain suggesting a task-modulating effect on S1 due to emotion processing.

**CONCLUSIONS:** Variation of daily life adaptive skills appears to be reflected in the brain’s mesoscale neural circuitry, as shown by the S1 connectivity profile, which is also differentially modulated during rest and emotional processing.

## Introduction

Sensory processing (SP) atypicalities constitute a prominent feature in the clinical presentation of autism spectrum disorders (ASD), affecting more than 90 per cent of the diagnosed individuals (Marco et al., 2011). ASD is primarily characterized by social, communication and/or restricted/repetitive behaviors (Lord et al., 2020), with SP differentiation only assessed as part of the latter according to the latest version of the Diagnostic and Statistical Manual of Mental Disorders (DSM-5).

Sensory processing difficulties are common across the ASD continuum. Therefore, there is a strong need to understand how these contribute to or interact with other core ASD symptoms. Some studies have previously shown that SP atypicalities are related to overall ASD severity -although more so at younger ages -as well as social difficulties (Kojovic et al., 2019), level of daily functioning (Suarez, 2012; Zachor & Ben-Itzchak, 2014), and adaptive behavior (Lane et al., 2010). Essentially, difficulties with sensory processing may amplify social interaction difficulties (Suarez, 2012), or be at their root. For instance, SP disruption has been found to be predictive of maladaptive behaviors both in cross-sectional (Baker et al., 2008; Lane et al., 2010), as well as in longitudinal studies (Williams et al., 2018). Moreover, tactile processing dysfunction has been explicitly linked to social difficulties in individuals with ASD (Hilton et al., 2010), highlighting the importance of touch in early development for forming relationships (Kropf et al., 2019). Despite the evidence suggesting a relationship between sensory processes and higher-order functions, such as those mentioned above, the neural underpinnings that could potentially link the two remain unclear.

In this work, we focus on the primary somatosensory cortex (S1), which is a core brain area for processing somatosensory information (Kropf et al., 2019; Mikkelsen et al., 2018). While mostly known for the processing of tactile information and pain (Bushnell et al., 1999; Kaas et al., 1979), growing evidence supports the notion that multisensory integration, which is essential for the perception of complex social information and that has been found to diverge in autistic individuals relative to neurotypicals (Marco et al., 2011), takes place within S1 and other sensory cortices as well (Kayser & Logothetis, 2007; Wallace et al., 2004). Furthermore, autistic adults appear to have a disrupted cortical representation of their face and hand, but there is a further need to understand the involvement of this region in ASD. For example, this area has been linked to emotional processing (Kropf et al., 2019) and it’s been suggested that lower-level tactile perception could be intact in autistic individuals, but differences in sensory under-or over-responsivity might be attributed to emotional difficulties instead (Mikkelsen et al., 2018).The link between higher-order cognitive variation (e.g., in emotional processing) and lower-order atypicalities of the somatosensory cortex deserves our attention and has been the target of this work.

Neurotypical or neurodivergent brain functioning of S1 can be understood and characterized through its topographic organization (Patel et al., 2014). This principle governs the S1 organization; adjacent parts of the body are represented in adjacent positions in the cortex. The somatotopic organization is typically being assessed through task-evoked activity where distinct areas of the body are being stimulated, resulting in distinct activation foci in S1 in alignment with what is known as sensory homunculus. A few methods have been proposed to directly capture the gradual nature of brain topographies; one of them is the ‘connectopic mapping’ approach (Haak et al., 2017). This method extends the primary approach of estimating connectivity between regions which have been defined based on hard parcellation schemes under the assumption of piece-wise constant connectivity patches and manages to capture smoothly-varying ‘connection topographies’/’connectopies’ from resting state fMRI (Haak et al., 2017, Marquand et al., 2017), without violating S1’s exhibited functional multiplicity (Haak & Beckmann, 2020). In previous work we established that the topographic map of primary somatosensory cortex closely maps on to the primary connectopic map within the motor complex of cortex M1/S1 (Haak et al., 2017). In the present work, S1 provides the ground for studying the neural underpinnings of low-level sensory processing, as well as their potential link to higher-order ASD features. Notably, such an examination ought to take place at an individual level, so that the mapping constructed between behavioral and biological scores takes into account the heterogeneity of the ASD condition, reported and discussed elsewhere (Zabihi et al., 2019), instead of relying on a case-control approach that could conceal such potential relationships (Marquand et al., 2016).

Here, we investigate: 1) whether S1’s topographic organization maps onto clinical scores measuring sensory processing, along with other daily life skills and abilities, and 2) how S1’s “rest topography” varies with emotional processing demands, in other words whether it is modulated by an emotion-eliciting task. To address these aims we applied connectopic mapping to the S1 cortex of neurotypical and autistic individuals in order to (i) find the association between the spatial characteristics of the estimated connection topographies and dimensional symptom scores across all individuals, and (ii) assess whether those spatial characteristics are different in an emotional processing task, relative to the resting state.

## Methods and Materials

### Neuroimaging data

The dataset that was used for the current analysis was part of the EU-AIMS LEAP project, a multi-site European collaborative effort with its main aim to identify biomarkers associated with ASD. The study design is described extensively in an earlier publication (Loth et al., 2017) along with a comprehensive overview of the clinical characterization of the cohort (Charman et al., 2017).

Resting-state functional MRI (rfMRI) scans were available for 656 participants, to which we applied a set of stringent quality control exclusion criteria. Participants with a structural brain abnormality (not clinically relevant; n=17), an incomplete rfMRI scan (n=8), excessive head motion during the rfMRI scan (mean frame-wise displacement>.7mm; n=29, max frame-wise displacement>3.8mm; n=38), or insufficient brain coverage (n=24) were excluded (n=145). We further excluded participants with insufficient variance of voxels in the selected region of interest (n=107; see following section *Connectopic Mapping for further details*). This resulted in the inclusion of 404 participants with rfMRI scans.

For the emotional processing task against which we compare our rest topographies, we used the Hariri emotion processing fMRI task (Hariri et al., 2002), for which 287 participants had data available. During this task participants were asked to decide which of the faces, displaying either a fearful or angry expression, or shapes presented to them at the top of screen matches the one presented at the bottom of the screen. After following the fMRI pre-processing and quality control (see Chauvin, 2019 for details), we included 249 participants in the analysis. Demographic and clinical characteristics of our sample for rfMRI and the Hariri task can be found in **Table *1***.

**Table 1.** Participant demographic and clinical characteristics (rfMRI, Hariri task).

|  | *rfMRI* |  | *Hariri Task* |  |
| --- | --- | --- | --- | --- |
|  | **N** | **Mean±SD** | **N** | **Mean±SD** |
| ***Demographic measures*** |  |  |  |  |
| **Neurotypical:autistic** | 164:240 | - | 125:124 | - |
| **Sex (female:male)** | 128:276 | - | 75:174 | - |
| **Handedness (left-handed:right-handed:ambidextrous) <sup>a</sup>** | 43:293:11 | - | 23:185:6 | - |
| **Age in years** | 404 | 16.74±5.3 | 249 | 17.49±5.48 |
| **Full-scale IQ** | 401 | 102.29±18.93 | 248 | 107.42±13.97 |
| ***Clinical measures*** |  |  |  |  |
| **SSP <sup>b</sup>** | 222 | 153.55±29.25 | 134 | 158.71±28.44 |
| **SRS-2 <sup>c</sup>** | 354 | 60.92±15.14 | 227 | 57.18±14.8 |
| **Vineland-II Communication domain** | 258 | 78.16±18.73 | 135 | 81.18±17.99 |
| **Vineland-II Daily living domain** | 256 | 76.82±18.66 | 134 | 80.15±17.47 |
| **Vineland-II Socialization domain** | 254 | 76.53±21.62 | 133 | 79.07±20.43 |
| **Vineland-II ABC** | 252 | 75.27±18.34 | 132 | 77.98±16.97 |
| **ADI-R Social domain** | 229 | 16.53±6.77 | 119 | 16.06±6.61 |
| **ADI-R Communication domain** | 228 | 13.37±5.57 | 118 | 13.03±5.62 |
| **ADI-R Restricted and Repetitive behaviors domain** | 217 | 4.5±2.44 | 112 | 4.28±2.61 |
| **ADOS-2 <sup>d</sup>** | 236 | 5.15±2.68 | 122 | 5.2±2.7 |

SD Standard Deviation, IQ Intelligence Quotient, SSP Short Sensory Profile, SRS-2 Social Responsiveness Scale-2, Vineland-II ABC Vineland-II Adaptive Behaviour Composite, ADI-R Autism Diagnostic Interview-Revised (only available for autistic individuals), ADOS-2 Autism Diagnostic Observation Schedule-Second Edition (only available for autistic individuals)
*a* Handedness information missing for 57 participants (rfMRI) and for 35 participants (Hariri).
*b* Total score (parent-report).
*c* Total T-score (combined parent- and self-report). Parent-report score prioritized over self-report score.
*d* Combine Calibrated Severity Scores (CSS) (Based on full data & imputed data).

### fMRI data acquisition

MRI data were acquired on different 3T scanners at multiple sites in Europe: General Electric MR750 (GE Medical Systems, Milwaukee, WI, USA) at Institute of Psychiatry, Psychology and Neuroscience, King’s College London, United Kingdom (KCL); Siemens Magnetom Skyra (Siemens, Erlangen, Germany) at Radboud University Nijmegen Medical Centre, the Netherlands (RUNMC); Siemens Magnetom Verio (Siemens, Erlangen, Germany) at the Autism Research Centre at the University of Cambridge, United Kingdom (UCAM); Philips 3T Achieva (Philips Healthcare Systems, Best, The Netherlands) at University Medical Centre Utrecht, the Netherlands (UMCU); and Siemens Magnetom Trio (Siemens, Erlangen, Germany) at Central Institute of Mental Health, Mannheim, Germany (CIMH) (see Supplementary Material). Structural images were obtained using a 5.5-minute MPRAGE sequence (TR=2300ms, TE=2.93ms, T1=900ms, voxels size=1.1×1.1×1.2mm, flip angle=9°, matrix size=256×256, FOV=270mm, 176 slices). An eight-to-ten minute resting-state fMRI (rfMRI) scan was acquired using a multi-echo planar imaging (ME-EPI) sequence; TR=2300ms, TE∼12ms, 31ms, and 48ms (slight variations are present across centers), flip angle=80°, matrix size=64×64, in-plane resolution=3.8mm, FOV=240mm, 33 axial slices, slice thickness/gap=3.8mm/0.4mm, volumes=200 (UMCU), 215 (KCL,CIMH), or 265 (RUNMC, UCAM). Participants were instructed to relax and fixate on a cross presented on the screen for the duration of the rfMRI scan (for further details see Oldehinkel et al., 2019).

### fMRI preprocessing

Preprocessing of the fMRI data was performed using a standard preprocessing pipeline that included tools from the FMRIB Software Library (FSL version 5.0.6; http://www.fmrib.ox.ac.uk/fsl). For the rfMRI data, we initially recombined the three rfMRI scan echoes using echo-time weighted averaging. Preprocessing included removal of the first five volumes to allow for signal equilibration, primary head motion correction via realignment to the middle volume using MCFLIRT (Jenkinson et al., 2002), global 4D mean intensity normalization and spatial smoothing with a 6mm FWHM Gaussian kernel. Then, ICA-AROMA was used to identify and remove secondary motion-related artifacts (Pruim et al., 2015; Pruim et al., 2015b). Next, nuisance regression was applied to remove signal from white matter and CSF, and a 0.01 Hz high-pass filter was applied to remove very low-frequency drifts in the time-series data. In order to preserve the broad-band characteristics of functional connectivity data no low-pass filtering was applied (Niazi, 2011). The fMRI images of each participant were co-registered to the participants’ high-resolution T1 anatomical images via boundary-based registration (BBR) implemented in FSL FLIRT (Jenkinson et al., 2002). The T1 images of each participant were registered to MNI152 standard space with FLIRT 12-dof linear registration (Andersson et al., 2007), and further refined using FNIRT non-linear registration (10mm warp, 2mm resampling resolution; Jenkinson et al., 2002). Finally, we brought all participant-level rfMRI images to 2mm MNI152 standard space, in which all further analyses were conducted.

### Clinical measures

The autistic participants have received clinical diagnosis according to the DSM-IV, ICD-10, or DSM-5 criteria. Furthermore, additional dimensional measures of sensory processing for both neurotypical and autistic participants were available. The *Short sensory profile* (SSP) (Tomchek & Dunn, 2007) was used to assess sensory processing atypicalities across 38 items from which an overall raw score was derived that reflects sensory processing across multiple sensory domains (lower scores indicate increased sensory atypicalities). The *Social Responsiveness Scale-Second Edition* (SRS-2, Bruni, 2014) was used to assess distinctions in social behavior associated with ASD across a 65-item rating scale. The overall T-scores from parent and self-reports (for neurotypical adults, SRS-2 self-report version of the questionnaire was administered, while for the rest a parent-reported symptom questionnaire was available) are reported here (higher scores suggest more severe social difficulties). Finally, the *Vineland Adaptive Behavior Scales, Second Edition* (Vineland-II; Sparrow et al., 2005) a semi-structured parent interview, was used to assess the adaptive functioning of participants across three domains: communication, socialization, and daily living skills (lower scores indicate lower adaptability skills in the respective life domain). For a more extensive description of all the existing clinical measures of this project, check (Charman et al., 2017). The differential availability of the measures led to a limited subgroup of participants examined for each measure (description of those measures can be found in **Table *1***). However, we repeated the analysis using imputed measures (missing values of autistic individuals were imputed using multivariate regression models; Llera et al., 2022), which led to similar conclusions.

### Connectopic Mapping

We selected the post-central gyrus based on the Harvard–Oxford atlas (available as part of FSL; Jenkinson et al., 2012) as our region of interest. Based on this ROI we estimated connection topographies (Haak et al., 2017) separately for each participant, hemisphere and imaging measure (rfMRI, Hariri task) using the *congrads* tool publicly available at https://github.com/koenhaak/congrads. The derived connectopies represent how the connectivity between the target ROI and the rest of neocortex vary topographically within the given ROI. All analyses were conducted in Montreal Neurological Institute 152 standard space.

We estimated a reference connectopy using 20 unrelated individuals from the Human Connectome Project (HCP) and used this as a reference for the comparison of the individual connectopies that we obtained in the LEAP dataset estimating the spatial correlation between them. The comparison between the connectopies of the two datasets will provide us with an estimation of the quality of the obtained connectopies, as well as of the validity of the main axis of connectivity change we observe, as HCP is a well-known excellent quality rfMRI dataset in which the somatotopic organization of the primary motor cortex – similar to that of the primary somatosensory cortex – has been mapped before (Haak et al., 2018). In order to sensitise our later analyses to subtle variations in this connectopy, individuals whose connectopies correlated less than r=0.5 with the reference HCP connectopy, together with the participants for whom connectopic mapping was not possible, were excluded from further analysis (n=107; see Section ‘Data preparation for connectopic mapping’ in Supplementary Material for analytic description of the criteria for exclusion and the steps followed).

Trend Surface Modelling (TSM) was applied on the connectopies providing us with a lower-dimensional representation to obtain a lower-dimensional representation which facilitates further statistical inference (Haak et al., 2017). By applying TSM on our individual connectopies, we obtain a set of parameters (TSM coefficients) that correspond to polynomial coefficients of varying order along 3D Euclidian axes. The polynomial degree, i.e the number of TSM coefficients, controls the granularity of the representation of the connectopic map and was chosen based on the value that minimized the model’s Bayesian information criterion (BIC) while, at the same time, maximized its explained variance (EV).

Additionally, we estimated the projection maps of the connectopies onto the rest of the brain by using dual regression, so as to map the whole-brain connectivity changes associated with the subtle variations within the ROI.

### Association with clinical measures

The Pearson’s correlation coefficient between the TSM coefficients of the connectopies, as generated according to the previous section and the available clinical measures was calculated. We additionally controlled for biological sex, age, full-scale IQ and site via OLS multiple regression. GLM analysis on raw connectopies was conducted using as explanatory variables ASD diagnosis, sex, age, and the clinical and behavioral scores reported in **Table *1***. P-values have been corrected via the Bonferroni-Holm method (FWER=0.05). We compared the connectopies between groups of neurotypical and autistic individuals, as well as between rest and task, using t-tests, independent and paired samples, respectively. The implementation code can be found at https://github.com/cisakoglou/congrads_sensory_leap.

## Results

### Average connectopies estimated across individuals during rest and the Hariri task

For the modeling of the S1 connectopy the 6^th^ order was selected for our model (see section ‘Model Selection’ in Supplementary Material). This entailed the representation of each connectopy with a set of 18 parameters (x, y, z, x^2^, y^2^, z^2^, x^3^, etc.) during both rest and task. That is, spatial variation within the ROI was modelled through a trend surface that was described by polynomials up to 6^th^ order in x,y and z direction.

The average connectopy across all individuals – autistic and neurotypical individuals combined – is illustrated in **Figure 1A** for rest and **Figure 1D** for task. After visual inspection, we observe the similarity with the average connectopy estimated from HCP (**Figure 1B**). The voxel-wise spatial correlation between the HCP connectopy and the one during each experimental condition was estimated (HCP-rest: (left) Pearson’s r=0.83, (right) Pearson’s r=0.94; HCP-task: (left) Pearson’s r=0.89, (right) Pearson’s r=0.96; p-values<.001 for all four comparisons).

The average S1 connectopy along the dorsoventral axis of each hemisphere was reproduced across both conditions and datasets with similar reconstructions of the connectopies for all (see reconstruction for the average S1 connectopy during rest in **Figure 1C**) and matches the somatotopic organization.

**Figure 1.**
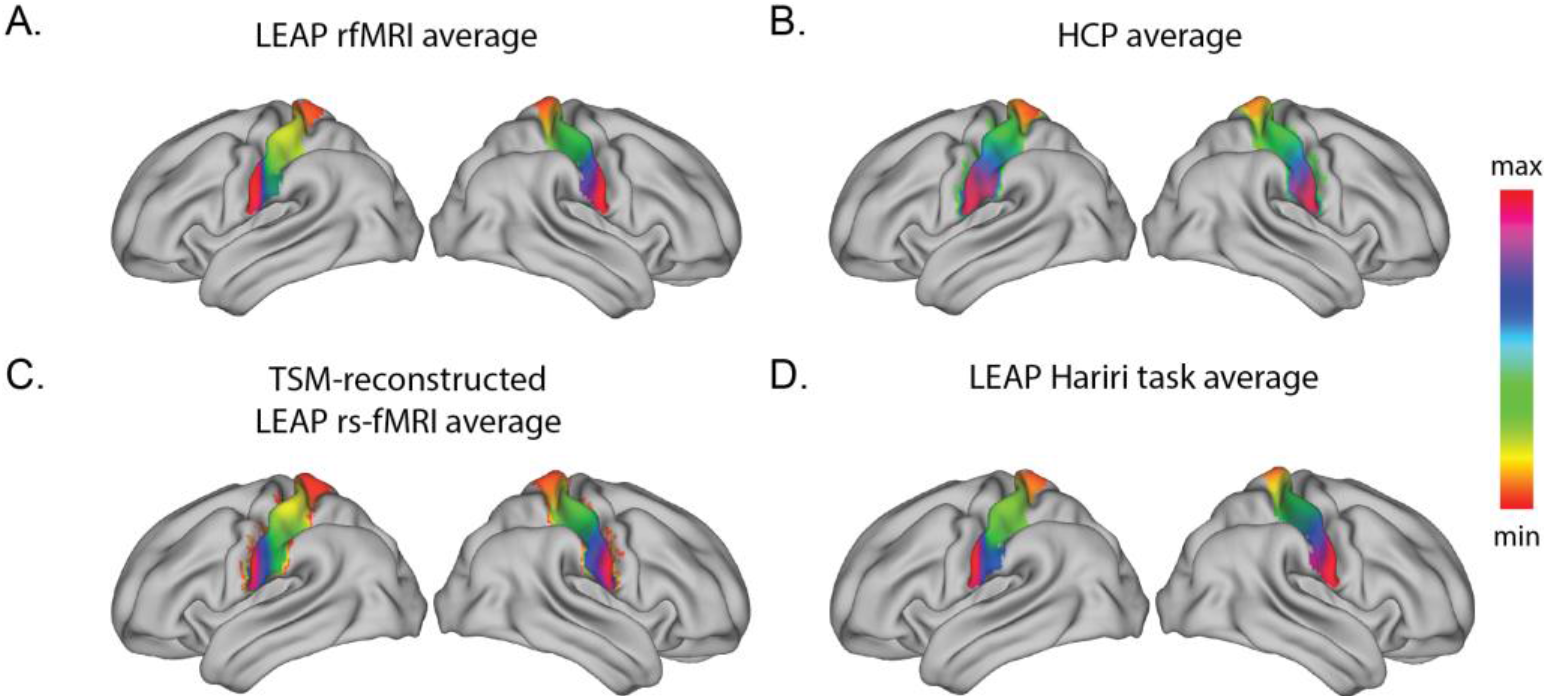
Primary somatosensory connectopies in LEAP and HCP datasets. Illustration for both hemispheres of the A. average primary connectopy of the LEAP rfMRI dataset after the application of connectopic mapping on the primary somatosensory cortex of N=404 participants, the B. HCP average S1 primary connectopy, used as a reference, the C. TSM-reconstructed gradient (6^th^ model order) of the LEAP rfMRI average primary connectopy (A), and the D. LEAP Hariri task average primary S1 connectopy. The connectopies for the two hemispheres were derived independently. The color bar indicates the position along the primary mode of connectivity change -similar colors represent similar connectivity patterns.

### Association between S1 connectopies and ASD dimensional measures

During the Hariri task, the association between two TSM coefficients (z^2^, z^4^) reconstructing the S1 connectopy on the left hemisphere and the Vineland-II Daily Living score values was found to be significant (see Supplementary Material). The correlations with Vineland-II Socialization and Adaptive Behavior Composite (ABC) standard scores were also high but did not survive multiple comparisons correction.

We conducted an additional GLM analysis on raw S1 connectopies to examine the localization of the above association. As reported in Supplementary Material, Vineland-II Daily-Living, Socialization and Communication scores survived correction and the association was mostly located at the borders between S1/M1.

During rest, some TSM coefficients (see Supplementary Material) were found to be correlated with the Vineland-II Daily Living and Vineland-II ABC standard scores. However, these correlations did not survive multiple comparisons correction. In this analysis, we controlled for chronological age, biological sex and acquisition site via confound regression. We also correlated TSM coefficients with SSP, SRS-2 and Vineland-II Socialization, and no significant association was identified.

We further aimed to understand how the S1 connectopy during the Hariri task changes as a function of the variation within the clinical measures. Having already obtained the TSM coefficients reconstructing the S1 connectopy and the respective clinical measure with which these were associated, we visualized the clinical associations of underlying topography in the following way; We selected three points across the functioning scale of the Vineland-II Daily Living score (1,2,3 as seen on ***Figure 2C***), we then derived their respective TSM-reconstructed connectopies (***Figure 2A***) by obtaining the residual connectopies after fitting their initial reconstructions to the average TSM-reconstructed connectopy (***Figure 2B***).

**Figure 2.**
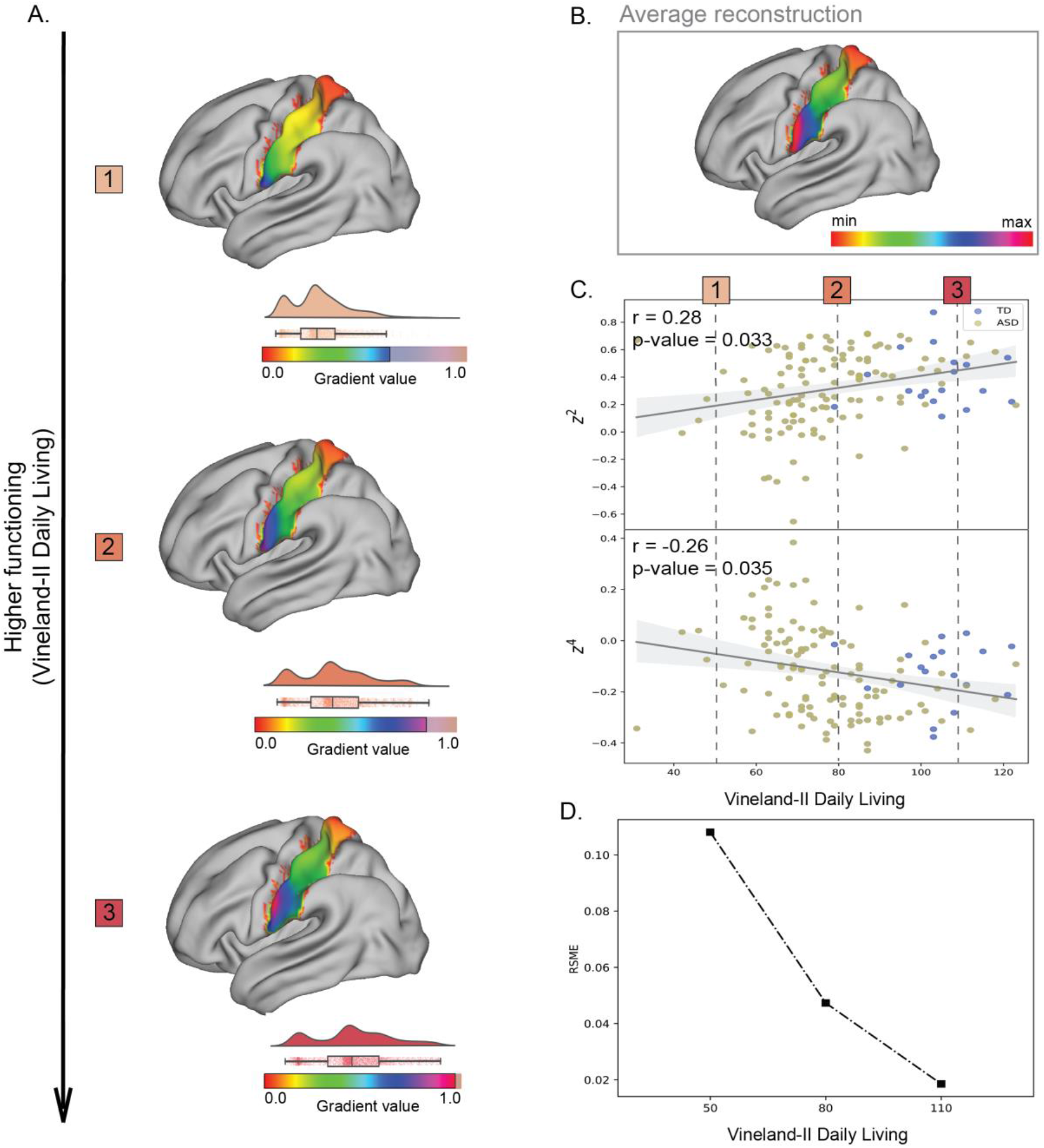
Visualization of the change in the connectivity profile within S1 of the left hemisphere during the Hariri task with respect to the functioning ability as reflected by Vineland-II Daily Living scores. A. The reconstructions of the connectopies corresponding to the three evenly spread points across the Vineland-II Daily Living scale, as indicated in C. In each of the three reconstructions, the gradient’s color range varies, and its variation is highlighted by a raincloud visualization of its gradient values, the B. TSM-reconstructed average S1 connectopy of the left hemisphere during the Hariri task, C. Scatterplots and correlation parameters between the TSM coefficients (6^th^ model order) of the S1 connectopies that were found to be significantly associated with Vineland-II Daily Living scores, and the D. RMSE between the reconstructed S1 connectopy of the each of the three selected points across Vineland-II Daily Living scale and the average S1 reconstructed connectopy.

We calculated the RMSE (root-mean-square error) between the TSM coefficients that reconstruct the average connectopy in the z direction – which includes the coefficients that were found to be correlated with the Vineland-II scores – and the TSM coefficients that reconstruct the connectopy corresponding to each of the selected points across the Vineland-II Daily Living scale that are highlighted in ***Figure 2C***. In ***Figure 2D***, we illustrated how the RMSE is reduced as a function of the adaptive Vineland-II scores, providing an estimation of the reduction of the flatness we observe within the S1 connectopy as functioning skills improve.

We showed that as adaptive functioning gets higher, which is expressed by the higher Vineland-II Daily Living scores, the color range within the gradient of the reconstructed connectopy becomes wider, reflecting a broader ROI to rest-of-brain differential connectivity pattern. Conversely, the gradient becomes flatter towards the other end of the functioning scale denoting lower functioning skills. The S1 connectopy of the individuals with higher adaptive functioning skills, spanning a broader range of differential connectivity along with the higher model order capturing the subtler changes of the connectopy, suggests higher functional differentiation in those individuals. Lower functional differentiation, on the other hand, is associated with lower adaptive functioning, but this does not indicate the absence of a specific function of S1.

### Comparison between the connectopies of neurotypical and autistic individuals

Group comparisons of the S1 connectopies between neurotypical and autistic individuals did not yield any significant results for either hemisphere for both experimental conditions. During the Hariri task, there were three TSM coefficients (y^2^, y^4^ and y^6^ [p-values: 0.017, 0.019, 0.025] for left hemisphere) that were nominally significant but did not survive multiple comparisons correction.

### Connectopic similarities and differences during rest and task

To elucidate further the nature of the established association between connectopies and adaptive scores, we examined in more detail the differences between intrinsic functional connectivity (iFC) gradients during rest and task connectopies. Despite the dorsoventral axis of the connectopy being reproduced in both rest and task, we observed significant differences between the two. The TSM coefficients that were found to differ in the 192 participants for whom the scans of both conditions were available were 6 coefficients on the z-axis for the left hemisphere and 5 coefficients on the x-and y-axis for the right one (see Supplementary Material). In

Figure 3 the differences between the rest and task connectopies are highlighted displaying not a focal, but rather a global pattern of differences along the dorsoventral axis of the S1.

**Figure 3.**
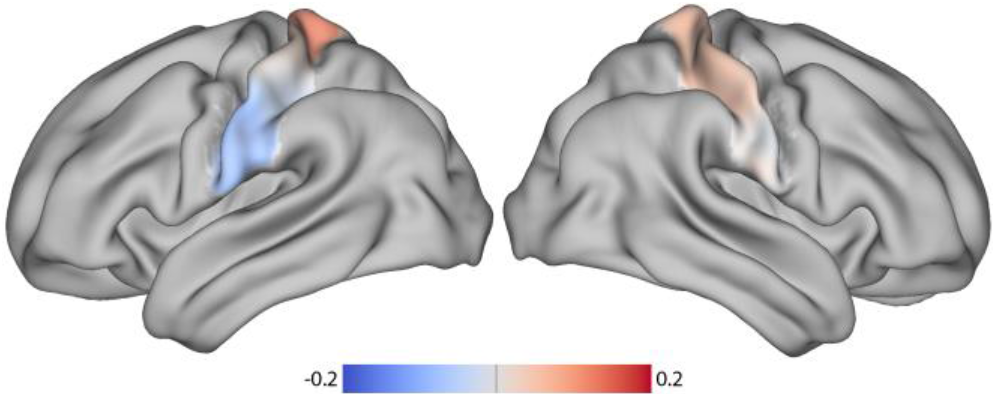
Differences between rfMRI and Hariri task primary connectopies. Visualization of the connectopic differences after averaging the differences between the individual connectopies during rfMRI and Hariri task (Connectopy_rfMRI_ – Connectopy_Hariri_).

We additionally estimated the projections of the S1 connectopies onto the brain in order to highlight further the spatial components involved in the connectivity profile changes we observe. Projections during rest from the left and right hemisphere’s S1 connectopy are illustrated respectively in **Figure 4A** and **Figure 4B** and for the Hariri task in **Figure 4C** and **Figure 4D**. Overall, the color-gradients present in the left/right hemisphere reappear contralaterally in the opposite hemisphere, providing evidence that the two sensory strips are topographically connected. There is a general convergence between the two experimental conditions (projections(left): Pearson’s r=0.85 [p-value<.001], projections(right): Pearson’s r=0.77 [p-value<.001]), apart from a frontal component – including anterior cingulate gyrus and frontal gyrus – that seem to have enhanced similarity in its connectivity profile with S1 during task and a prefrontal component, on the opposing end, with a highlighted lack of similarity during task, more evident for the right hemisphere.

**Figure 4.**
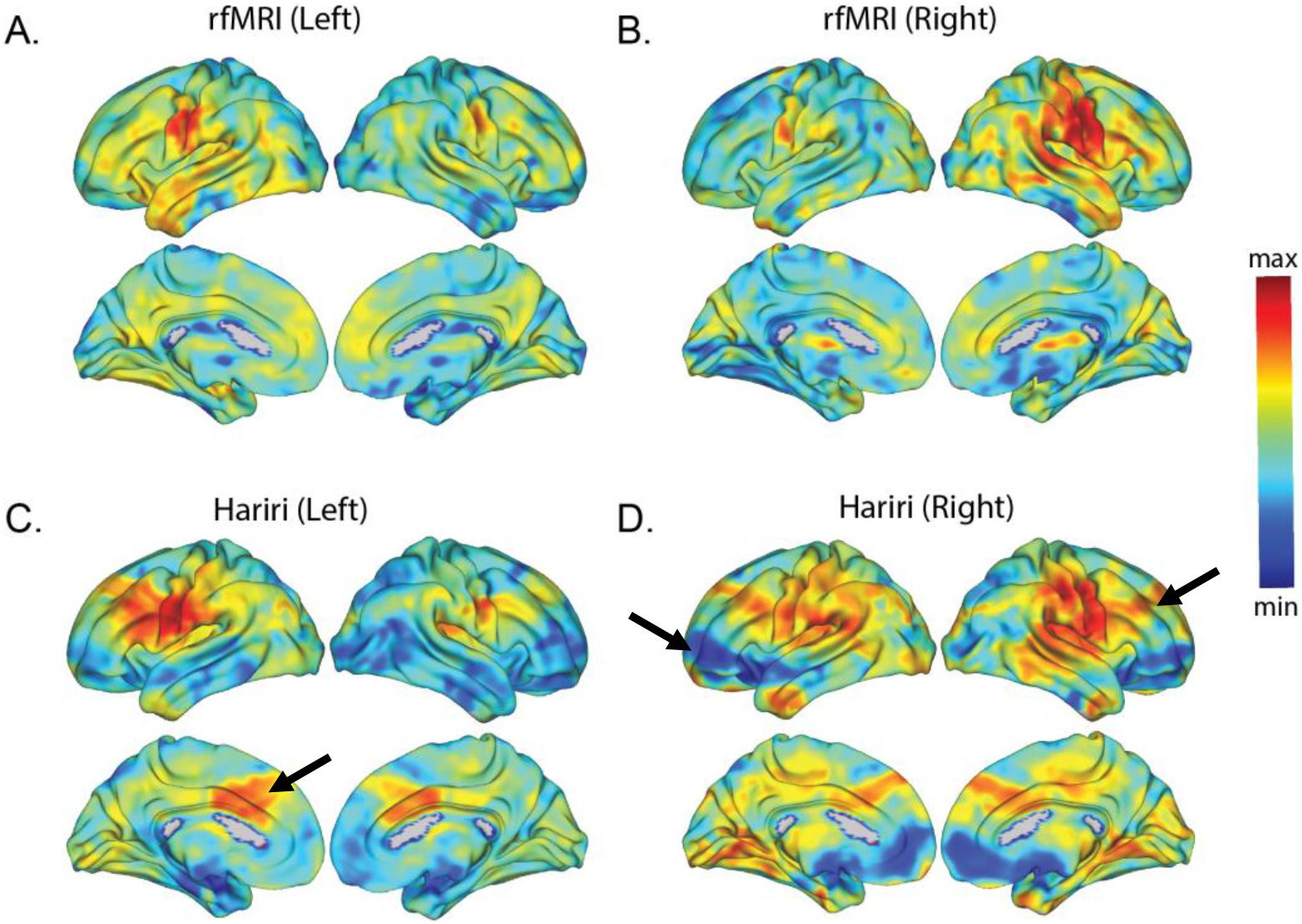
Projection maps. Projection maps of the S1 connectopies onto the entire brain during the rfMRI from the left and right hemisphere in A and B, respectively, and during the Hariri task from the left and right hemisphere in C and D, respectively.

## Discussion

In this study, we investigated in a data-driven manner the fine-grained connectivity of the primary somatosensory cortex in the context of ASD and its potential cascading effects on key ASD traits. We detected an association between S1 connectopy during the emotional matching task and participants’ adaptive abilities in relation to their daily lives. We showed how S1 connectopy reflects variation in adaptive abilities, illustrating a gradient that follows the well-established somatotopic organization of a dorsomedial-to-ventrolateral axis (i.e., homunculus) in the left hemisphere of individuals and gradually flattens as their daily life adaptive abilities decline. We found no case-control differences between neurotypical and autistic individuals on either their Ifc gradients or task gradients. Last, we examined the differences in S1 connectopies at rest and during task. The differences found there, present both in the S1 connectopies themselves and in their projection maps, led us to postulate that S1 is also influenced in a top-down manner, and in a way that its changes are then associated with permanent individual traits.

Our findings provide evidence that underlying biological mechanisms involved in lower-order sensory processing map onto higher-order behavioral atypicalities in ASD at the individual level. This extends recent discussions about the interplay between sensory atypicalities, such as under-responsiveness to touch and active seeking of sensational experiences, and social and communicative ones, as well as adaptive behavior (Lane et al., 2010; Mikkelsen et al., 2018; Suarez, 2012). In our work, we established a link with the individuals’ ability to adapt in their daily lives, as assessed by the Vineland-II Daily Living scale. An individual’s assessment on this scale is associated with real-word outcomes, such as the individual’s likelihood of independent living and social competence, and thus, attains great importance for what affects autistic individuals in their day-to-day matters. The cascading effects of an aberrant connectivity profile within the S1 region as illustrated here builds upon the idea suggested elsewhere (Marco et al., 2011a) that potential disruptions in unimodal sensory information processing are forming the backbone of higher order cortical abilities.

Notably, in this work we did not establish an association between S1 connectivity profile and sensory processing atypicalities, as reflected by the parent-report SSP scores. This could be due to a variety of reasons, from which the potential inadequacy of the TSM coefficients to capture effectively the gradual connectopy was excluded, as our GLM analysis on the raw connectopies replicated the same relationships as the ones revealed using the TSM coefficients. Our work is also susceptible to the potential limitations of SSP scores, such as the small number of items, and quiet disperse over the different sensory modalities (tactile atypicalities corresponding only to a fraction of the questionnaire’s items), that may stand unable to capture the wide heterogeneity of sensory phenotypes present in ASD (Tillmann et al., 2020). Adding to this, the wide age range included in this work may perplex things further when taking into consideration the developmental component of ASD sensory phenotype, limiting even further the power of the current sample, and as an extension our ability, to capture any potentially relevant relationship. Of note, studies so far on the neurobiology of sensory profiles report using only a subset of SSP items and Sensory over-responsivity scores (Green et al., 2015), as it was argued that while SSP is beneficial for clinical image formulation, it can show less correlation with brain structure than direct assessment (Tavassoli et al., 2019).

Our estimated S1 connectopies were following, as it would be expected, a dorsomedial-to-ventrolateral axis that matches the established somatotopic organization of the region (Patel et al., 2014; Roux et al., 2018; Yeo et al., 2011). However, by visualizing in more detail the profile of connectivity changes within the S1 region at specific points on the Vineland-II Daily Living scale we were able to show that the lower the adaptive skills of the individual, the less functionally differentiated the gradient describing them. On the contrary, the better an individual functions in daily life, the broader the color range of their S1 gradients appears to become, which translated to higher variability of connectivity profiles within this region and potentially stronger differentiation in the associated pattern of connectivity. Our results align with and build upon a growing body of evidence suggesting that ASD occurs due to altered communication between brain regions, and most specifically due to reduced functional segregation of large-scale brain networks (Hong et al., 2019; Rudie et al., 2012). A recent study, in which authors utilized a global connectopic mapping approach, has furthermore shown that the segregation between sensory systems and unimodal, as well as transmodal, convergence regions appears to be diminished in ASD (Hong et al., 2019). In our work, we further bind the notion of varying degree of functional differentiation within S1 with a varying degree of the adaptive behavior in the daily life of individuals.

Adding to the discussion of how and whether the focus on the average patient may in fact disguise inter-individual differences in psychiatry (Wolfers et al., 2019), the lack of case-control differences in our results alludes to the known heterogeneity of ASD (Zabihi et al., 2019b) and underscores the need of looking into individual variations instead. Despite the disrupted cortical connectivity theory being one of the prominent ones with regards to ASD and its etiology, the localization and identification of the exact mechanisms (i.e., what kind of connectivity disruption, its behavioral link) involved remain elusive so far (Rane et al., 2015). On top of the heterogeneity characterizing ASD, potentially being the main reason behind the aforementioned elusiveness of findings, lies the assumption of piece-wise constant connectivity that most studies have been so far relying on and that neglects the functional heterogeneity (continuous change) with respect to connectivity of certain regions (Haak & Beckmann, 2020). Nevertheless, what our work highlights is the need for a description that takes place at the individual level, characterizes connectivity change in a gradual fashion and examines its association with core ASD phenotypes.

We further established topographic differences in S1 between rest and the Hariri emotional matching paradigm. This provides evidence for a disrupted cortical hierarchy due to higher-order processes taking place during emotion recognition which would require the presence of non-unilateral connections in S1. Differences in the brain regions recruited by autistic individuals for the purposes of multisensory integration in the context of emotion recognition have been found before (Thye et al., 2018), but our results further highlight the direct involvement of S1 into this interplay and could be beneficial for conditions characterized by co-occurring emotional and sensory difficulties. Additionally, the existence of a frontal component in the projection maps of the S1 connectopies present during task, and not during rest, support the idea that S1 is not solely responsible for the low-level processing of sensory stimuli. The frontal component is mostly located around the area of cingulate gyrus that has been found to have great influence on social behavior (Apps et al., 2016), as well as on regulation of emotional processing (Posner et al., 2007), and has been found to be involved in both the sensory and social atypicalities observed in ASD (Thye et al., 2018). Evidence for the functional role of S1 in perceiving emotional categories has been reported before (Kragel & LaBar, 2016), while emotional face processing has been repeatedly linked to ASD severity (Hadjikhani et al., 2004). All this, along with the association between daily living adaptive skills and within S1 topographic variation, puts S1 on the map of the sensory-social axis in ASD, and provides evidence for social and adaptive functioning in daily life being dependent on perceiving and recognizing others’ emotions, and this on its own being dependent on sensory processing.

All of the above taken together, suggest a potential overarching link between higher- and lower-level processes in the brain, relying on long-range bottom-up, as well as top-down, connections, whose aberrant communication will be reflected on someone’s daily life functioning skills. Our findings are consistent with studies that mention atypical sensory features in ASD being a consequence of both bottom-up, as well as top-down processing distinctions (Green et al., 2015) and highlight a prospective transdiagnostic value for S1 in conditions where sensory distress co-occurs with higher-order cognitive atypicalities.

### Limitations

Although we provide convincing evidence for the validity of the S1 connectopies using two individual datasets and experimental conditions, there is a certain limitation that needs to be highlighted. We use the Harvard-Oxford anatomical atlas, which, although well established in our field, remains a probabilistic atlas that does not take into account the functional role of the selected areas. This could lead to signal contamination that interferes with the alignment between the actual functional boundaries of the selected region for S1 and the boundaries used in this work.

### Future directions

In future the investigation of the output from connectopic mapping by its application on distinct sensory modalities, e.g., auditory, face processing of the somatosensory – in the latter case higher spatial resolution would be required. Additionally, the discrepancy between rest and task could be quantified further and compared against ASD severity and other dimensional scores. Another important aspect of the sensory atypicalities that we cannot address by solely looking at a single time point is their developmental effect. This could be investigated further by making use of the subsequent timepoint data present in the LEAP dataset. The assessment of inter-individual differences in a paired design, rather than a cross-sectional analysis could prove more sensitive with respect to the effects studied here, and even though we had the same individuals available for two experimental conditions, this could be extended for multiple timepoints. Lastly, as sensory hyper-and hypo-responsivity is not unique to ASD (but is more common in this population than in other developmental disorders), the relationship examined in the current work could be explored in other clinical populations.

## Conclusion

In this study, we showed that individual variation present in the dorsoventral connectopies within the primary somatosensory cortex is linked to ASD dimensional measures that access the daily life adaptive skills of individuals. Furthermore, differences between the primary connectopies and their projection maps during rest and task suggest a modulating role of the primary somatosensory cortex within a possible top-down regulation of low-level somatosensory processing framework. The findings of this study lay the foundation for the generation of more specific hypotheses regarding the mechanisms of sensory processing dysfunction in ASD and provide an opportunity for earlier phenotypic features that can help with the diagnosis of ASD.

## Acknowledgments

This work is primarily supported by the EU-AIMS consortium (European Autism Interventions), which receives support from Innovative Medicines Initiative Joint Undertaking Grant No.115300, the resources of which are composed of financial contributions from the European Union’s Seventh Framework Programme (Grant No. FP7/2007-2013), from the European Federation of Pharmaceutical Industries and Associations companies’ in-kind contributions; and by the AIMS-2-TRIALS consortium (Autism Innovative Medicine Studies-2-Trials), which has received funding from the Innovative Medicines Initiative 2 Joint Undertaking under grant agreement No. 777394, and this Joint Undertaking receives support from the European Union’s Horizon 2020 research and innovation programme and EFPIA and AUTISM SPEAKS, Autistica, SFARI. This work reflects the authors’ views and neither IMI nor the European Union, EFPIA or any Associated Partners are responsible for any use that may be made of the information contained therein.

CI is supported by funding from the Gravitation Programme ‘Language in Interaction’ Grant No. 024.001.006 from the Netherlands Organization for Scientific Research. KVH gratefully acknowledges funding from the Dutch Research Council (Veni 016.171.068 and Vidi 09150171910043). TW gratefully acknowledges the European Union’s Horizon 2020 research and innovation programme under the Marie Sklodowska-Curie grant agreement No 895011. DLF is supported by funding from the European Union’s Horizon 2020 research and innovation programme under the Marie Skłodowska-Curie grant agreement No 101025785. AL was supported by CANDY (Grant No. 847818). JKB was supported by the Horizon2020 supported programme CANDY (Grant No. 847818). AFM gratefully acknowledges support from the Netherlands Institute for Scientific research under a VIDI grant (016.156.415) and an ERC consolidator grant (10100118). CFB gratefully acknowledges funding from the Wellcome Trust Collaborative Award in Science 215573/Z/19/Z and the Netherlands Organization for Scientific Research Vici Grant No. 17854 and NWO-CAS Grant No. 012-200-013.

## References

Andersson, J. L. R., Jenkinson, M., Smith, S., & others. (2007). Non-linear registration, aka Spatial normalisation FMRIB technical report TR07JA2. FMRIB Analysis Group of the University of Oxford, 2(1), e21.

Apps, M. A. J., Rushworth, M. F. S., & Chang, S. W. C. (2016). The Anterior Cingulate Gyrus and Social Cognition: Tracking the Motivation of Others. In Neuron (Vol. 90, Issue 4, pp. 692–707). Cell Press. https://doi.org/10.1016/j.neuron.2016.04.018

Baker, A. E. Z., Lane, A., Angley, M. T., & Young, R. L. (2008). The relationship between sensory processing patterns and behavioural responsiveness in autistic disorder: A pilot study. Journal of Autism and Developmental Disorders, 38(5), 867–875. https://doi.org/10.1007/s10803-007-0459-0

Bruni, T. P. (2014). Test Review: Social Responsiveness Scale–Second Edition (SRS-2). Journal of Psychoeducational Assessment, 32(4), 365–369. https://doi.org/10.1177/0734282913517525

Bushnell, M. C., Duncan, G. H., Hofbauer, R. K., Ha, B., Chen, J. I., & Carrier, B. (1999). Pain perception: Is there a role for primary somatosensory cortex? Proceedings of the National Academy of Sciences of the United States of America, 96(14), 7705–7709. https://doi.org/10.1073/pnas.96.14.7705

Charman, T., Loth, E., Tillmann, J., Crawley, D., Wooldridge, C., Goyard, D., Ahmad, J., Auyeung, B., Ambrosino, S., Banaschewski, T., Baron-Cohen, S., Baumeister, S., Beckmann, C., Bölte, S., Bourgeron, T., Bours, C., Brammer, M., Brandeis, D., Brogna, C., … Buitelaar, J. K. (2017). The EU-AIMS Longitudinal European Autism Project (LEAP): Clinical characterisation. Molecular Autism, 8(1), 1–21. https://doi.org/10.1186/s13229-017-0145-9

Chauvin, R. J. M. (2019). The efficient brain. On how connectivity modulations underpin cognitive tasks (Doctoral dissertation) [Radboud University]. https://hdl.handle.net/2066/210159

Green, S. A., Hernandez, L., Tottenham, N., Krasileva, K., Bookheimer, S. Y., & Dapretto, M. (2015). Neurobiology of sensory overresponsivity in youth with autism spectrum disorders. JAMA Psychiatry, 72(8), 778–786. https://doi.org/10.1001/jamapsychiatry.2015.0737

Haak, K. V., & Beckmann, C. F. (2020). Understanding brain organisation in the face of functional heterogeneity and functional multiplicity. NeuroImage, 220, 117061. https://doi.org/10.1016/j.neuroimage.2020.117061

Haak, K. V., Marquand, A. F., & Beckmann, C. F. (2017). Connectopic mapping with resting-state fMRI. NeuroImage, 170, 83–94. https://doi.org/10.1016/j.neuroimage.2017.06.075

Haak, K. V., Marquand, A. F., & Beckmann, C. F. (2018). Connectopic mapping with resting-state fMRI. In NeuroImage (Vol. 170, pp. 83–94). Academic Press Inc. https://doi.org/10.1016/j.neuroimage.2017.06.075

Hadjikhani, N., Joseph, R. M., Snyder, J., Chabris, C. F., Clark, J., Steele, S., McGrath, L., Vangel, M., Aharon, I., Feczko, E., Harris, G. J., & Tager-Flusberg, H. (2004). Activation of the fusiform gyrus when individuals with autism spectrum disorder view faces. NeuroImage, 22(3), 1141–1150. https://doi.org/10.1016/j.neuroimage.2004.03.025

Hariri, A. R., Tessitore, A., Mattay, V. S., Fera, F., & Weinberger, D. R. (2002). The amygdala response to emotional stimuli: A comparison of faces and scenes. NeuroImage, 17(1), 317–323. https://doi.org/10.1006/nimg.2002.1179

Hilton, C. L., Harper, J. D., Kueker, R. H., Lang, A. R., Abbacchi, A. M., Todorov, A., & Lavesser, P. D. (2010). Sensory responsiveness as a predictor of social severity in children with high functioning autism spectrum disorders. Journal of Autism and Developmental Disorders, 40(8), 937–945. https://doi.org/10.1007/s10803-010-0944-8

Hong, S. J., de Wael, R. V., Bethlehem, R. A. I., Lariviere, S., Paquola, C., Valk, S. L., Milham, M. P., Di Martino, A., Margulies, D. S., Smallwood, J., & Bernhardt, B. C. (2019). Atypical functional connectome hierarchy in autism. Nature Communications, 10(1), 1–13. https://doi.org/10.1038/s41467-019-08944-1

Jenkinson, M., Bannister, P., Brady, M., & Smith, S. (2002). Improved optimization for the robust and accurate linear registration and motion correction of brain images. NeuroImage, 17(2), 825–841. https://doi.org/10.1016/S1053-8119(02)91132-8

Jenkinson, M., Beckmann, C. F., Behrens, T. E. J., Woolrich, M. W., & Smith, S. M. (2012). FSL. NeuroImage, 62(2), 782–790. https://doi.org/10.1016/j.neuroimage.2011.09.015

Kaas, J. H., Nelson, R. J., Sur, M., Lin, C. S., & Merzenich, M. M. (1979). Multiple representations of the body within the primary somatosensory cortex of primates. Science, 204(4392), 521–523. https://doi.org/10.1126/science.107591

Kayser, C., & Logothetis, N. K. (2007). Do early sensory cortices integrate cross-modal information? In Brain Structure and Function (Vol. 212, Issue 2, pp. 121–132). Springer. https://doi.org/10.1007/s00429-007-0154-0

Kojovic, Ben Hadid Franchini, & Schaer. (2019). Sensory Processing Issues and Their Association with Social Difficulties in Children with Autism Spectrum Disorders. Journal of Clinical Medicine, 8(10), 1508. https://doi.org/10.3390/jcm8101508

Kragel, P. A., & LaBar, K. S. (2016). Somatosensory representations link the perception of emotional expressions and sensory experience. ENeuro, 3(2), 169–177. https://doi.org/10.1523/ENEURO.0090-15.2016

Kropf, E., Syan, S. K., Minuzzi, L., & Frey, B. N. (2019). From anatomy to function: the role of the somatosensory cortex in emotional regulation. Revista Brasileira de Psiquiatria (Sao Paulo, Brazil : 1999), 41(3), 261–269. https://doi.org/10.1590/1516-4446-2018-0183

Lane, A. E., Young, R. L., Baker, A. E. Z., & Angley, M. T. (2010). Sensory processing subtypes in autism: Association with adaptive behavior. Journal of Autism and Developmental Disorders, 40(1), 112–122. https://doi.org/10.1007/s10803-009-0840-2

Llera, A., Brammer, M., Oakley, B., Tillmann, J., Zabihi, M., Mei, T., Charman, T., Ecker, C., Acqua, F. D., Banaschewski, T., Moessnang, C., Baron-Cohen, S., Holt, R., Durston, S., Murphy, D., Loth, E., Buitelaar, J. K., Floris, D. L., & Beckmann, C. F. (2022). Evaluation of data imputation strategies in complex, deeply-phenotyped data sets: the case of the EU-AIMS Longitudinal European Autism Project. http://arxiv.org/abs/2201.09753

Lord, C., Brugha, T. S., Charman, T., Cusack, J., Dumas, G., Frazier, T., Jones, E. J. H., Jones, R. M., Pickles, A., State, M. W., Taylor, J. L., & Veenstra-VanderWeele, J. (2020). Autism spectrum disorder. Nature Reviews Disease Primers, 6(1). https://doi.org/10.1038/s41572-019-0138-4

Loth, E., Charman, T., Mason, L., Tillmann, J., Jones, E. J. H., Wooldridge, C., Ahmad, J., Auyeung, B., Brogna, C., Ambrosino, S., Banaschewski, T., Baron-Cohen, S., Baumeister, S., Beckmann, C., Brammer, M., Brandeis, D., Bölte, S., Bourgeron, T., Bours, C., … Buitelaar, J. K. (2017). The EU-AIMS Longitudinal European Autism Project (LEAP): Design and methodologies to identify and validate stratification biomarkers for autism spectrum disorders. Molecular Autism, 8(1), 1–19. https://doi.org/10.1186/s13229-017-0146-8

Marco, E. J., Hinkley, L. B. N., Hill, S. S., & Nagarajan, S. S. (2011a). Sensory Processing in Autism: A Review of Neurophysiologic Findings. Pediatric Research, 69(5 Pt 2), 48R–54R. https://doi.org/10.1203/PDR.0b013e3182130c54

Marco, E. J., Hinkley, L. B. N., Hill, S. S., & Nagarajan, S. S. (2011b). Sensory Processing in Autism: A Review of Neurophysiologic Findings. Pediatric Research, 69(5 Pt 2), 48R–54R. https://doi.org/10.1203/PDR.0b013e3182130c54

Marquand, A. F., Haak, K. V., & Beckmann, C. F. (2017). Functional corticostriatal connection topographies predict goal-directed behaviour in humans. Nature Human Behaviour, 1(8), 1–23. https://doi.org/10.1038/s41562-017-0146

Marquand, A. F., Rezek, I., Buitelaar, J., & Beckmann, C. F. (2016). Understanding Heterogeneity in Clinical Cohorts Using Normative Models: Beyond Case-Control Studies. Biological Psychiatry, 80(7), 552–561. https://doi.org/10.1016/j.biopsych.2015.12.023

Mikkelsen, M., Wodka, E. L., Mostofsky, S. H., & Puts, N. A. J. (2018). Autism spectrum disorder in the scope of tactile processing. Developmental Cognitive Neuroscience, 29, 140–150. https://doi.org/10.1016/j.dcn.2016.12.005

Niazi, M. K. K. (2011). Image Filtering Methods for Biomedical Applications. Acta Universitatis Upsaliensis.

Oldehinkel, M., Mennes, M., Marquand, A., Charman, T., Tillmann, J., Ecker, C., Dell’Acqua, F., Brandeis, D., Banaschewski, T., Baumeister, S., Moessnang, C., Baron-Cohen, S., Holt, R., Bölte, S., Durston, S., Kundu, P., Lombardo, M. V., Spooren, W., Loth, E., … Zwiers, M. P. (2019). Altered Connectivity Between Cerebellum, Visual, and Sensory-Motor Networks in Autism Spectrum Disorder: Results from the EU-AIMS Longitudinal European Autism Project. Biological Psychiatry: Cognitive Neuroscience and Neuroimaging, 4(3), 260–270. https://doi.org/10.1016/j.bpsc.2018.11.010

Patel, G. H., Michael, D. K., & Snyder, L. H. (2014). Topographic organization in the brain: Searching for general principles. Trends in Cognitive Sciences, 18(7), 351–363. https://doi.org/10.1038/jid.2014.371

Posner, M. I., Rothbart, M. K., Sheese, B. E., & Tang, Y. (2007). The anterior cingulate gyrus and the mechanism of self-regulation. In Cognitive, Affective and Behavioral Neuroscience (Vol. 7, Issue 4, pp. 391–395). Springer. https://doi.org/10.3758/CABN.7.4.391

Pruim, R. H. R., Mennes, M., Buitelaar, J. K., & Beckmann, C. F. (2015). Evaluation of ICA-AROMA and alternative strategies for motion artifact removal in resting state fMRI. NeuroImage, 112, 278–287. https://doi.org/10.1016/j.neuroimage.2015.02.063

Pruim, R. H. R., Mennes, M., van Rooij, D., Llera, A., Buitelaar, J. K., & Beckmann, C. F. (2015). ICA-AROMA: A robust ICA-based strategy for removing motion artifacts from fMRI data. NeuroImage, 112, 267–277. https://doi.org/10.1016/j.neuroimage.2015.02.064

Rane, P., Cochran, D., Hodge, S. M., Haselgrove, C., Kennedy, D. N., & Frazier, J. A. (2015). Connectivity in Autism: A Review of MRI Connectivity Studies. In Harvard Review of Psychiatry (Vol. 23, Issue 4, pp. 223–244). Taylor and Francis Ltd. https://doi.org/10.1097/HRP.0000000000000072

Roux, F.-E., Djidjeli, I., & Durand, J.-B. (2018). Functional architecture of the somatosensory homunculus detected by electrostimulation. The Journal of Physiology, 596(5), 941–956. https://doi.org/10.1113/JP275243

Rudie, J. D., Shehzad, Z., Hernandez, L. M., Colich, N. L., Bookheimer, S. Y., Iacoboni, M., & Dapretto, M. (2012). Reduced functional integration and segregation of distributed neural systems underlying social and emotional information processing in Autism spectrum disorders. Cerebral Cortex, 22(5), 1025–1037. https://doi.org/10.1093/cercor/bhr171

Sparrow, S. S., Cicchetti, D. V, & Balla, D. A. (2005). Vineland adaptive behavior scales:(Vineland II), survey interview form/caregiver rating form. Livonia, MN: Pearson Assessments.

Suarez, M. A. (2012). Sensory processing in children with autism spectrum disorders and impact on functioning. In Pediatric Clinics of North America (Vol. 59, Issue 1, pp. 203–214). Elsevier. https://doi.org/10.1016/j.pcl.2011.10.012

Tavassoli, T., Brandes-Aitken, A., Chu, R., Porter, L., Schoen, S., Miller, L. J., Gerdes, M. R., Owen, J., Mukherjee, P., & Marco, E. J. (2019). Sensory over-responsivity: parent report, direct assessment measures, and neural architecture. Molecular Autism, 10(1), 4. https://doi.org/10.1186/s13229-019-0255-7

Thye, M. D., Bednarz, H. M., Herringshaw, A. J., Sartin, E. B., & Kana, R. K. (2018). The impact of atypical sensory processing on social impairments in autism spectrum disorder. Developmental Cognitive Neuroscience, 29(May 2017), 151–167. https://doi.org/10.1016/j.dcn.2017.04.010

Tillmann, J., Uljarevic, M., Crawley, D., Dumas, G., Loth, E., Murphy, D., Buitelaar, J., Charman, T., Ahmad, J., Ambrosino, S., Auyeung, B., Baumeister, S., Beckmann, C., Bourgeron, T., Bours, C., Brammer, M., Brandeis, D., Brogna, C., De Bruijn, Y., … Zwiers, M. P. (2020). Dissecting the phenotypic heterogeneity in sensory features in autism spectrum disorder: A factor mixture modelling approach. Molecular Autism, 11(1), 67. https://doi.org/10.1186/s13229-020-00367-w

Tomchek, S. D., & Dunn, W. (2007). Sensory processing in children with and without autism: a comparative study using the short sensory profile. American Journal of Occupational Therapy, 61(2), 190–200. https://doi.org/10.5014/ajot.61.2.190

Wallace, M. T., Ramachandran, R., & Stein, B. E. (2004). A revised view of sensory cortical parcellation. Proceedings of the National Academy of Sciences of the United States of America, 101(7), 2167–2172. https://doi.org/10.1073/pnas.0305697101

Williams, K. L., Kirby, A. V., Watson, L. R., Sideris, J., Bulluck, J., & Baranek, G. T. (2018). Sensory features as predictors of adaptive behaviors: A comparative longitudinal study of children with autism spectrum disorder and other developmental disabilities. Research in Developmental Disabilities, 81, 103–112. https://doi.org/10.1016/j.ridd.2018.07.002

Wolfers, T., Beckmann, C. F., Hoogman, M., Buitelaar, J. K., Franke, B., & Marquand, A. F. (2019). Individual differences v. the average patient: Mapping the heterogeneity in ADHD using normative models. Psychological Medicine, 50(2), 314–323. https://doi.org/10.1017/S0033291719000084

Yeo, B. T. T., Krienen, F. M., Sepulcre, J., Sabuncu, M. R., Lashkari, D., Hollinshead, M., Roffman, J. L., Smoller, J. W., Zollei, L., Polimeni, J. R., Fischl, B., Liu, H., & Buckner, R. L. (2011). The organization of the human cerebral cortex estimated by intrinsic functional connectivity. Journal of Neurophysiology, 106, 1125–1165. https://doi.org/10.1152/jn.00338.2011.

Zabihi, M., Oldehinkel, M., Wolfers, T., Frouin, V., Goyard, D., Loth, E., Charman, T., Tillmann, J., Banaschewski, T., Dumas, G., Holt, R., Baron-Cohen, S., Durston, S., Bölte, S., Murphy, D., Ecker, C., Buitelaar, J. K., Beckmann, C. F., & Marquand, A. F. (2019a). Dissecting the Heterogeneous Cortical Anatomy of Autism Spectrum Disorder Using Normative Models. Biological Psychiatry: Cognitive Neuroscience and Neuroimaging, 4(6), 567–578. https://doi.org/10.1016/j.bpsc.2018.11.013

Zabihi, M., Oldehinkel, M., Wolfers, T., Frouin, V., Goyard, D., Loth, E., Charman, T., Tillmann, J., Banaschewski, T., Dumas, G., Holt, R., Baron-Cohen, S., Durston, S., Bölte, S., Murphy, D., Ecker, C., Buitelaar, J. K., Beckmann, C. F., & Marquand, A. F. (2019b). Dissecting the Heterogeneous Cortical Anatomy of Autism Spectrum Disorder Using Normative Models. Biological Psychiatry: Cognitive Neuroscience and Neuroimaging, 4(6), 567–578. https://doi.org/10.1016/j.bpsc.2018.11.013

Zachor, D. A., & Ben-Itzchak, E. (2014). The relationship between clinical presentation and unusual sensory interests in autism spectrum disorders: A preliminary investigation. Journal of Autism and Developmental Disorders, 44(1), 229–235. https://doi.org/10.1007/s10803-013-1867-y

