## Supplementary Material for "Fine-grained topographic organization within somatosensory cortex during resting-state and emotional face-matching task and its association with ASD traits"

### Methods

#### *Data preparation for connectopic mapping*

For rs-fMRI, we excluded the participants for which there were more than 5000 voxels with invalid values (inf/nan) within the region of our interest (n=8). We then shrank the ROI by removing the voxels in which there was at least one participant that had invalid values and used this as the common roi for estimating all individual connectopies. We further excluded the participants whose connectopies had spatial correlation with the HCP reference connectopy less than 0.5 (n=99). For the Hariri task, we used the ROI we obtained during rs-fMRI and we excluded the participants that had any missing value within it (n=38). One extra participant was also excluded due to missing clinical information.

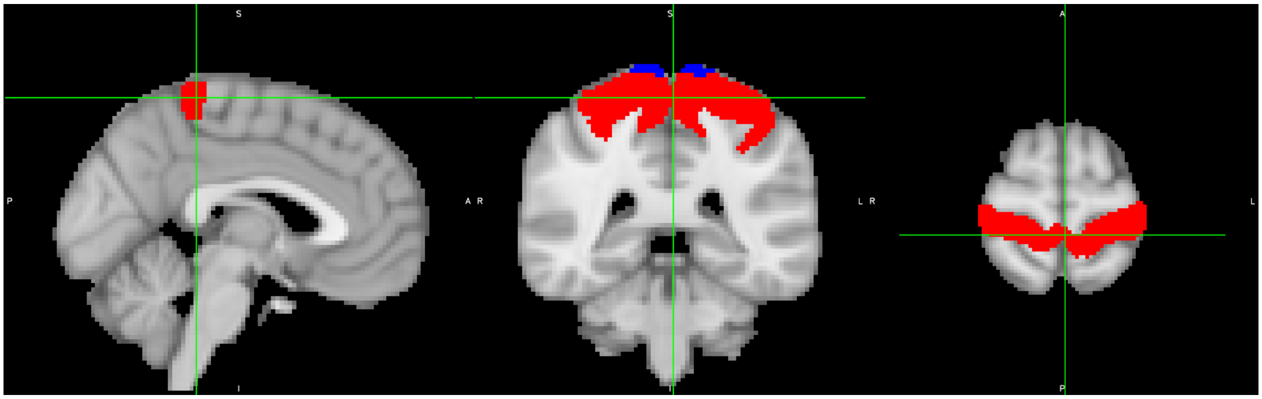

*Figure 1 Obtained ROI after the procedure described above is displayed in red on top of the original ROI. The blue part seen here is the part for which there were invalid values for at least one participant, and hence removed.*

*Sample characteristics*

| | N | ALL (N=404) | AUTISTIC (N=240) | NEUROTYPICAL (N=164) |
|---|---|---|---|---|
| Autistic:Neurotypical | 404 | 240:164 | | |
| Gender (f:m) | | 128:276 | 62:178 | 66:98 |
| Age in years [mean (SD)] | | 16.74 [5.3] | 16.95 [5.25] | 16.43 [5.37] |
| Full-scale IQ [mean (SD)] | 401 | 102.29 [18.93] | 100.08 [19.03] | 105.5 [18.37] |
| Handedness (l:r:a) | 347 | 43:293:11 | 29:173:8 | 14:120:3 |
| *Clinical measures* | | | | |
| SSP [mean (SD)] | 222 | 153.55 [29.25] | 139.03 [26.73] | 177.86 [12.19] |
| SRS-2 [mean (SD)] | 354 | 60.92 [15.14] | 69.74 [11.82] | 46.76 [6.9] |
| Vineland-II [mean (SD)] Communication domain | 258 | 78.16 [18.73] | 75.03 [15.34] | 89.96 [24.91] |
| Vineland-II [mean (SD)] Daily living domain | 258 | 76.82 [18.66] | 73.27 [15.88] | 90.11 [22.16] |
| Vineland-II [mean (SD)] Socialization domain | 254 | 76.53 [21.62] | 71.09 [16.48] | 97.15 [26.14] |
| Vineland-II ABC | 252 | 75.27 [18.34] | 71.02 [13.53] | 91.26 [24.42] |
| ADI-R [mean (SD)] (Social domain) | 229 | - | 16.53 [6.77] | - |
| ADI-R [mean (SD)] (Communication domain) | 228 | - | 13.37 [5.57] | - |
| ADI-R [mean (SD)] (Restricted and Repetitive behaviours domain) | 217 | - | 4.5 [2.44] | - |
| ADOS-2 [l] | 236 | - | 5.15 [2.68] | - |

*Table 1 Demographics and clinical characteristics for the participants included during rs-fMRI.*

| | N | ALL (N=249) | AUTISTIC (N=124) | NEUROTYPICAL (N=125) |
|---|---|---|---|---|
| Autistic:Neurotypical | 249 | 124:125 | | |
| Gender (f:m) | | 75:174 | 39:85 | 36:89 |
| Age in years [mean (SD)] | 249 | 17.49 [5.48] | 17.79 [5.7] | 17.19 [5.26] |
| Full-scale IQ [mean (SD)] | 248 | 107.42 [13.97] | 107.06 [15.9] | 107.78 [11.78] |
| Handedness (l:r:a) | 214 | 23:185:6 | 11:94:3 | 12:91:3 |
| *Clinical measures* | | | | |
| SSP [mean (SD)] | 134 | 158.71 [28.44] | 143.45 [27.87] | 178.71 [12.16] |
| SRS-2 [mean (SD)] | 227 | 57.18 [14.8] | 67.56 [13.16] | 46.52 [6.47] |
| Vineland-II [mean (SD)] Communication domain | 135 | 81.18 [17.99] | 77.37 [15.79] | 105.94 [9.97] |
| Vineland-II [mean (SD)] Daily living domain | 134 | 80.15 [17.47] | 76.43 [15.24] | 104.11 [10.68] |
| Vineland-II [mean (SD)] Socialization domain | 133 | 79.07 [20.43] | 73.81 [16.07] | 112.67 [11.28] |
| Vineland-II ABC | 132 | 77.98[16.97] | 73.52 [13.27] | 106.22 [8.35] |
| ADI-R [mean (SD)] (Social domain) | 119 | - | 16.06 [6.61] | - |
| ADI-R [mean (SD)] (Communication domain) | 118 | - | 13.03 [5.62] | - |
| ADI-R [mean (SD)] | 112 | - | 4.28 [2.61] | - |

| | | | | |
|---|---|---|---|---|
| (Restricted and Repetitive behaviours domain) | | | | |
| ADOS-2 | 122 | - | 5.2 [2.7] | - |

*Table 2 Demographics and clinical characteristics for the participants included during the Hariri task.*

SD Standard Deviation, IQ Intelligence Quotient, SSP Short Sensory Profile, SRS-2 Social Responsiveness Scale-2, Vineland-II ABC Vineland-II Adaptive Behaviour Composite, ADI-R Autism Diagnostic Interview-Revised, ADOS-2 Autism Diagnostic Observation Schedule-Second Edition

### Results

#### *Reconstruction of average connectopy for neurotypical individuals during the Hariri task*

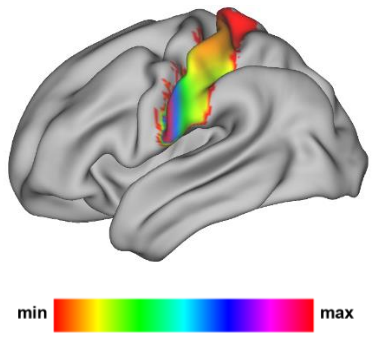

*Figure 2 Reconstruction of average connectopy for neurotypical individuals for which the Vineland-II Daily Living scores are available during the Hariri task (left hemisphere).*

*Model Selection*

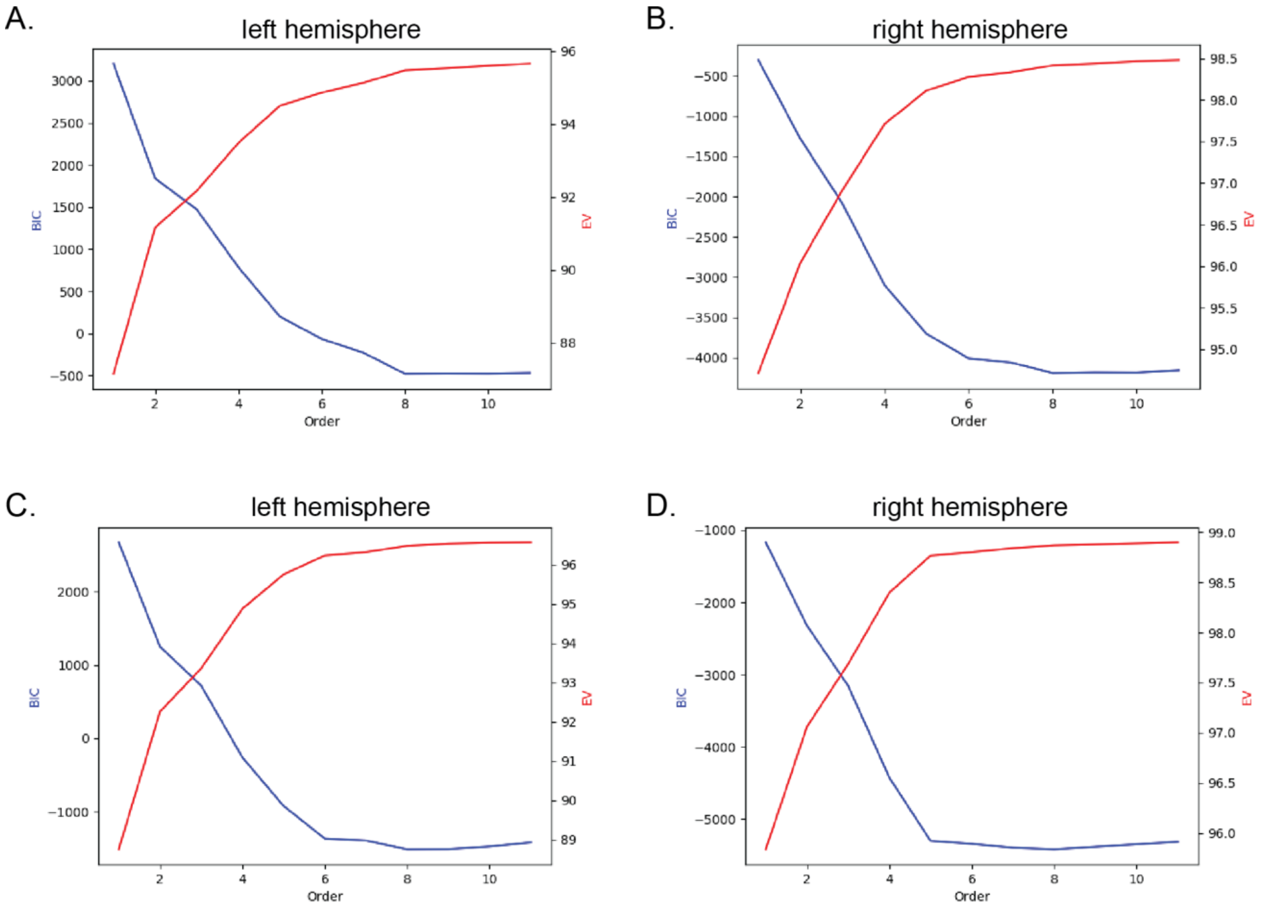

*Figure 3* ***Model order selection process.*** *The model order that simultaneously minimizes BIC and maximizes EV was selected. A. and B. show the BIC and EV as a function of model order during rs-fMRI for the left and right hemispheres, respectively, while C. and D. show the same quantities for the task condition and for the left and right hemispheres, respectively. Above the 6th model order, the model was not drastically improved for all four conditions. Therefore, in order not to increase the complexity of our model without achieving a large benefit that would justify its use, we decided to use this model order.*

#### *Association between assessment scores and spatial coefficients of connectopies*

| Scores | x | y | z | x2 | y2 | z2 | x3 | y3 | z3 | x4 | y4 | z4 | x5 | y5 | z5 | x6 | y6 | z6 |
|---|---|---|---|---|---|---|---|---|---|---|---|---|---|---|---|---|---|---|
| **SRS-2** | -0.03 | 0.07 | 0.02 | 0.02 | 0.11 | -0.08 | -0.03 | -0.04 | 0.06 | -0.02 | -0.09 | 0.05 | 0.06 | 0.04 | -0.04 | -0.03 | 0.08 | -0.05 |
| **SSP** | 0.02 | -0.16 | 0.03 | -0.11 | -0.08 | 0.21 | 0.06 | 0.16 | -0.07 | 0.02 | 0.06 | -0.17 | -0.1 | -0.16 | 0.03 | 0.07 | -0.05 | 0.11 |
| **RBS-R** | 0 | 0.14 | 0.03 | 0.01 | -0.03 | -0.14 | -0.03 | -0.12 | -0.03 | 0.05 | 0.05 | 0.15 | 0.06 | 0.12 | 0.07 | -0.09 | -0.05 | -0.07 |
| **ADI-R Social** | 0.14 | 0.19 | -0.1 | 0.26 | 0.03 | -0.19 | -0.07 | -0.18 | 0.1 | -0.2 | -0.04 | 0.21 | 0.01 | 0.17 | -0.1 | 0.12 | 0.04 | -0.21 |
| **ADI-R Comm.** | 0.15 | 0.27 | -0.09 | 0.11 | 0.06 | -0.14 | -0.03 | -0.11 | 0.14 | -0.1 | -0.02 | 0.17 | -0.02 | 0.1 | -0.15 | 0.08 | 0.01 | -0.21 |
| **ADI-R RBS** | 0.12 | 0.23 | -0.04 | 0.12 | 0.06 | -0.14 | -0.05 | -0.17 | 0.06 | -0.11 | -0.05 | 0.18 | 0 | 0.2 | -0.05 | 0.08 | 0.04 | -0.17 |
| **Vineland-II Comm.** | 0 | -0.12 | 0.01 | -0.09 | -0.1 | 0.14 | 0.02 | 0.14 | -0.12 | 0.1 | 0.08 | -0.11 | -0.01 | -0.1 | 0.12 | -0.07 | -0.08 | 0.12 |
| **Vineland-II D.Liv.** | -0.02 | -0.18 | 0.11 | -0.18 | -0.05 | **0.28*** | -0.03 | 0.17 | -0.16 | 0.14 | 0.04 | **-0.26*** | 0.06 | -0.14 | 0.11 | -0.12 | -0.04 | 0.21 |
| **Vineland-II Soc.** | 0.07 | -0.12 | 0.13 | -0.12 | -0.06 | 0.24 | -0.12 | 0.14 | -0.22 | 0.11 | 0.05 | -0.23 | 0.1 | -0.13 | 0.18 | -0.1 | -0.04 | 0.24 |
| **Vineland-II ABC** | 0.02 | -0.15 | 0.08 | -0.18 | -0.08 | 0.23 | -0.02 | 0.18 | -0.19 | 0.14 | 0.09 | -0.2 | 0.02 | -0.15 | 0.18 | -0.09 | -0.09 | 0.23 |

*Figure 4* ***Correlation analysis between the TSM parameters of the S1 primary connectopies during the Hariri task and the ASD clinical/behavioral scores for the left hemisphere.*** *Heatmap correlations [*: used to denote statistical significance after controlling and correcting multiple comparisons correction, SRS-2: Social Responsiveness Scale-2, RBS-R: Repetitive Behaviors Scale-Revised, ADI-R: Autism Diagnostic Interview-Revised, Vineland-II Comm.: Vineland-II Communication, Vineland-II D.Liv.: Vineland-II Daily Living, Vineland-II Soc.: Vineland-II Socialisation, Vineland-II-ABC: Vineland-II Adaptive Behaviour Composite (ABC) standard score.*

| Scores | x | y | z | x2 | y2 | z2 | x3 | y3 | z3 | x4 | y4 | z4 | x5 | y5 | z5 | x6 | y6 | z6 |
|---|---|---|---|---|---|---|---|---|---|---|---|---|---|---|---|---|---|---|
| **SRS-2** | 0.03 | 0.09 | 0.05 | 0.03 | 0.04 | 0.01 | -0.09 | 0.07 | 0.01 | -0.05 | -0.06 | -0.04 | 0.08 | -0.11 | -0.02 | 0.06 | 0.07 | 0.02 |
| **SSP** | -0.01 | -0.18 | -0.05 | -0.02 | 0.01 | -0.06 | 0.05 | 0.03 | -0.11 | 0.05 | 0.01 | 0.03 | -0.06 | 0.01 | 0.11 | -0.08 | -0.01 | 0.03 |
| **RBS-R** | -0.03 | 0.03 | 0.03 | -0.07 | -0.01 | 0.12 | 0.02 | 0.08 | 0 | 0.05 | 0 | -0.1 | -0.02 | -0.13 | 0 | -0.04 | 0 | 0.07 |
| **ADI-R Social** | -0.05 | 0.03 | 0.09 | -0.09 | 0.15 | 0.03 | 0.12 | 0.08 | -0.02 | 0.08 | -0.13 | -0.06 | -0.14 | -0.11 | 0.02 | -0.09 | 0.12 | 0.06 |
| **ADI-R Comm.** | 0.06 | 0.09 | 0.05 | -0.16 | 0.08 | 0.11 | -0.02 | 0.01 | 0.03 | 0.12 | -0.05 | -0.07 | -0.03 | -0.03 | 0 | -0.11 | 0.03 | 0.06 |
| **ADI-R RBS** | -0.06 | 0.1 | 0.06 | 0.05 | 0.01 | 0.11 | 0.06 | -0.04 | 0.07 | -0.05 | -0.03 | -0.09 | -0.04 | -0.01 | -0.11 | 0.04 | 0.02 | -0.02 |
| **Vineland-II Comm.** | -0.05 | 0.18 | -0.09 | -0.17 | -0.19 | 0.09 | -0.01 | -0.2 | -0.01 | 0.16 | 0.24 | 0.04 | -0.01 | 0.25 | 0.09 | -0.12 | -0.25 | 0.03 |
| **Vineland-II D.Liv.** | -0.02 | 0.09 | -0.01 | 0.02 | -0.16 | 0.12 | -0.04 | -0.19 | -0.06 | -0.01 | 0.15 | -0.06 | 0.05 | 0.26 | 0.08 | 0.01 | -0.15 | 0.07 |
| **Vineland-II Soc.** | -0.01 | 0.06 | 0.02 | -0.06 | -0.04 | 0 | -0.02 | -0.1 | -0.06 | 0.05 | 0.07 | 0 | 0.02 | 0.14 | 0.06 | -0.04 | -0.08 | 0.03 |
| **Vineland-II ABC** | -0.04 | 0.1 | -0.04 | -0.07 | -0.14 | 0.09 | -0.02 | -0.14 | -0.04 | 0.07 | 0.17 | -0.03 | 0.02 | 0.21 | 0.04 | -0.05 | -0.17 | 0.02 |

*Figure 5* ***Correlation analysis between the TSM parameters of the S1 primary connectopies during the Hariri task and the ASD clinical/behavioral scores for the right hemisphere.*** *Heatmap correlations [*: used to denote statistical significance after controlling and correcting multiple comparisons correction, SRS-2: Social Responsiveness Scale-2, RBS-R: Repetitive Behaviors Scale-Revised, ADI-R: Autism Diagnostic Interview-Revised, Vineland-II Comm.: Vineland-II Communication, Vineland-II D.Liv.: Vineland-II Daily Living, Vineland-II Soc.: Vineland-II Socialisation, Vineland-II-ABC: Vineland-II Adaptive Behaviour Composite (ABC) standard score*

| Scores | x | y | z | x2 | y2 | z2 | x3 | y3 | z3 | x4 | y4 | z4 | x5 | y5 | z5 | x6 | y6 | z6 |
|---|---|---|---|---|---|---|---|---|---|---|---|---|---|---|---|---|---|---|
| SRS-2 | 0.03 | 0.08 | 0.05 | 0.03 | 0.09 | -0.03 | -0.03 | -0.05 | 0.01 | -0.03 | -0.1 | 0.01 | 0.02 | 0.04 | -0.04 | 0.02 | 0.1 | -0.02 |
| SSP | -0.01 | -0.08 | -0.08 | 0 | -0.06 | 0.06 | -0.01 | 0.07 | 0.04 | 0.01 | 0.07 | -0.04 | 0.03 | -0.05 | -0.04 | -0.03 | -0.07 | 0 |
| RBS-R | 0.07 | 0.15 | 0.04 | -0.05 | 0.11 | -0.06 | -0.03 | -0.07 | -0.01 | 0.05 | -0.07 | 0.02 | 0 | 0.04 | 0 | -0.01 | 0.05 | -0.01 |
| ADI-R Social | 0.06 | 0 | -0.01 | 0.1 | -0.09 | 0 | -0.06 | -0.07 | -0.09 | -0.03 | 0.04 | -0.01 | 0.04 | 0.08 | 0.11 | -0.01 | -0.02 | 0.06 |
| ADI-R Comm. | 0.05 | 0.06 | -0.08 | 0.14 | 0.01 | -0.07 | -0.06 | -0.12 | 0.04 | -0.06 | -0.04 | 0.05 | 0.04 | 0.09 | -0.01 | 0.01 | 0.04 | -0.02 |
| ADI-R RBS | 0 | 0.11 | 0.04 | 0.03 | 0.14 | -0.01 | -0.11 | -0.2 | -0.02 | 0 | -0.17 | -0.03 | 0.1 | 0.18 | 0.02 | -0.05 | 0.16 | 0.04 |
| Vineland-II Comm. | 0.09 | 0.12 | -0.09 | -0.17 | -0.08 | -0.02 | 0.01 | 0.03 | 0 | 0.15 | 0.16 | 0.06 | -0.03 | -0.05 | 0.05 | -0.07 | -0.17 | -0.01 |
| Vineland-II D.Liv. | 0.09 | 0.1 | -0.13 | -0.2 | -0.12 | -0.08 | 0.06 | 0.08 | 0.05 | 0.16 | 0.23 | 0.12 | -0.08 | -0.08 | 0.01 | -0.04 | -0.25 | -0.06 |
| Vineland-II Soc. | 0.01 | 0.07 | -0.11 | -0.13 | -0.08 | -0.06 | 0.08 | 0.01 | 0.06 | 0.09 | 0.16 | 0.1 | -0.09 | -0.02 | -0.02 | -0.01 | -0.16 | -0.06 |
| Vineland-II ABC | 0.07 | 0.11 | -0.13 | -0.17 | -0.1 | -0.06 | 0.06 | 0.03 | 0.05 | 0.14 | 0.2 | 0.11 | -0.08 | -0.04 | 0.01 | -0.03 | -0.21 | -0.06 |

*Figure 6* ***Correlation analysis between the TSM parameters of the S1 primary connectopies during rs-fMRI and the ASD clinical/behavioral scores for the left hemisphere.*** *Heatmap correlations [*: used to denote statistical significance after controlling and correcting multiple comparisons correction, SRS-2: Social Responsiveness Scale-2, RBS-R: Repetitive Behaviors Scale-Revised, ADI-R: Autism Diagnostic Interview-Revised, Vineland-II Comm.: Vineland-II Communication, Vineland-II D.Liv.: Vineland-II Daily Living, Vineland-II Soc.: Vineland-II Socialisation, Vineland-II-ABC: Vineland-II Adaptive Behaviour Composite (ABC) standard score.*

| Scores | x | y | z | x2 | y2 | z2 | x3 | y3 | z3 | x4 | y4 | z4 | x5 | y5 | z5 | x6 | y6 | z6 |
|---|---|---|---|---|---|---|---|---|---|---|---|---|---|---|---|---|---|---|
| SRS-2 | -0.01 | 0.04 | 0.09 | 0 | 0.07 | 0.04 | 0 | 0.05 | -0.05 | -0.02 | -0.08 | -0.07 | 0.01 | -0.09 | 0.04 | 0.02 | 0.07 | 0.07 |
| SSP | 0.02 | -0.03 | -0.12 | 0 | -0.04 | -0.04 | 0 | -0.05 | 0.02 | 0.04 | 0.05 | 0.03 | -0.02 | 0.07 | 0 | -0.04 | -0.04 | 0 |
| RBS-R | -0.04 | 0.02 | 0.06 | 0.02 | 0.04 | 0.08 | 0.06 | -0.02 | 0.01 | -0.05 | -0.03 | -0.05 | -0.03 | 0.03 | 0 | 0.05 | 0.03 | 0.03 |
| ADI-R Social | -0.06 | 0.08 | -0.03 | 0.04 | -0.03 | -0.01 | 0.04 | -0.11 | -0.04 | 0.01 | 0.01 | 0.01 | -0.03 | 0.13 | 0.07 | -0.02 | -0.02 | 0.04 |
| ADI-R Comm. | -0.06 | 0.11 | -0.08 | 0.1 | 0 | -0.07 | 0.03 | -0.11 | 0.03 | -0.03 | -0.01 | 0.07 | -0.02 | 0.09 | -0.01 | 0.01 | 0.01 | -0.04 |
| ADI-R RBS | 0.01 | 0.1 | 0.03 | 0.04 | 0.13 | 0.01 | 0.03 | -0.14 | 0.02 | -0.04 | -0.13 | -0.04 | -0.03 | 0.11 | -0.03 | 0.02 | 0.12 | 0 |
| Vineland-II Comm. | -0.05 | -0.04 | -0.03 | -0.16 | -0.1 | 0.05 | 0.02 | 0.07 | -0.05 | 0.1 | 0.15 | 0.01 | -0.01 | -0.02 | 0.07 | -0.07 | -0.15 | 0.04 |
| Vineland-II D.Liv. | -0.04 | -0.05 | 0 | -0.2 | -0.09 | 0.04 | -0.02 | 0.15 | -0.06 | 0.12 | 0.16 | 0 | 0.03 | -0.11 | 0.07 | -0.07 | -0.16 | 0.04 |
| Vineland-II Soc. | -0.02 | -0.03 | -0.03 | -0.18 | -0.05 | -0.02 | -0.02 | 0.07 | -0.03 | 0.12 | 0.11 | 0.06 | 0.02 | -0.03 | 0.05 | -0.08 | -0.12 | 0.01 |
| Vineland-II ABC | -0.04 | -0.05 | -0.03 | -0.19 | -0.09 | 0.03 | -0.01 | 0.1 | -0.04 | 0.13 | 0.16 | 0.03 | 0.01 | -0.05 | 0.07 | -0.08 | -0.16 | 0.03 |

*Figure 7* ***Correlation analysis between the TSM parameters of the S1 primary connectopies during rs-fMRI and the ASD clinical/behavioral scores for the right hemisphere.*** *Heatmap correlations [*: used to denote statistical significance after controlling and correcting multiple comparisons correction, SRS-2: Social Responsiveness Scale-2, RBS-R: Repetitive Behaviors Scale-Revised, ADI-R: Autism Diagnostic Interview-Revised, Vineland-II Comm.: Vineland-II Communication, Vineland-II D.Liv.: Vineland-II Daily Living, Vineland-II Soc.: Vineland-II Socialisation, Vineland-II-ABC: Vineland-II Adaptive Behaviour Composite (ABC) standard score.*

*GLM analysis on raw connectopies during the Hariri task*

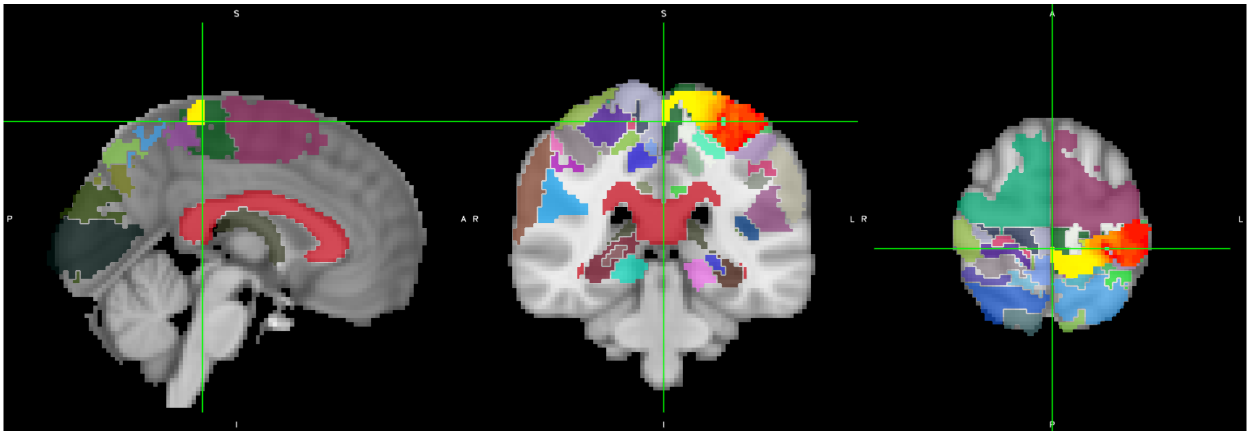

*Figure 8 T-maps for Vineland-II Daily living scores, superimposed on the Juelich atlas*

*(Only the significant voxels after correction are shown in the red-yellow color range).*

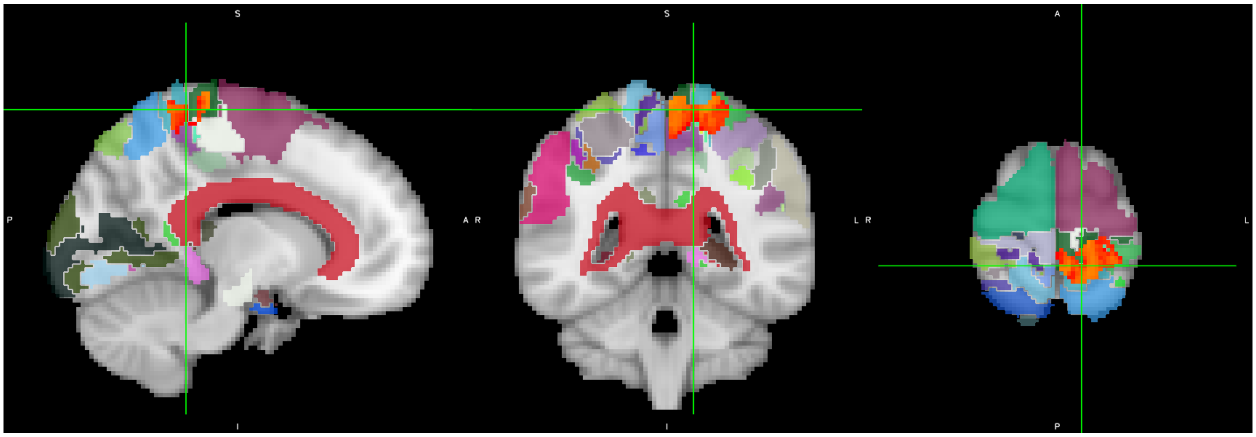

*Figure 9 T-maps for Vineland-II Socialization scores, superimposed on the Juelich atlas*

*(Only the significant voxels after correction are shown in the red-yellow color range).*

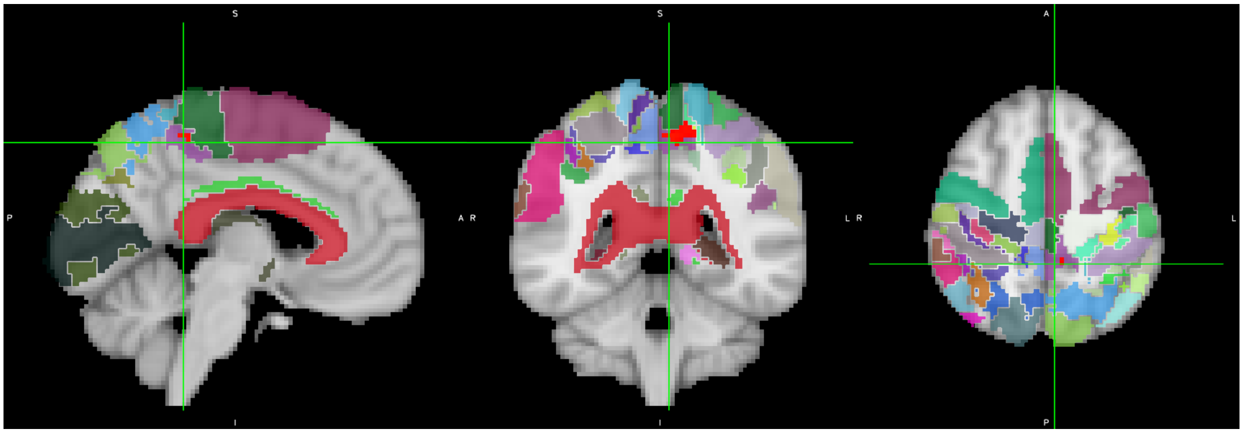

*Figure 10 T-maps for Vineland-II Communication scores, superimposed on the Juelich atlas*

*(Only the significant voxels after correction are shown in the red-yellow color range).*

##### *Differences between rfMRI and Hariri task connectopies*

TSM parameters that were found to differ between rest and task connectopies for left and right hemisphere, along with their respective corrected p-values.

| Hemisphere | Left | Right |
|---|---|---|
| **TSM parameters** | $z$, $z^2$, $z^3$, $z^4$, $z^5$, $z^6$ | $x^2$, $x^4$, $y^4$, $x^6$, $y^6$ |
| **p-values** | 0.0011, 0.0053, 0.0003, 0.0002, 0.0075, 0.0002 | 3e-06, 0.0001, 0.0073, 0.0028, 0.0038 |

#### Implementation

For the implementation of all the statistical tests, the Python module statsmodels (v0.9.0) and scipy.stats (v1.2.1) was used (group comparison between neurotypical and Autistic individuals s performed using ttest_ind, and group comparison between rest and Hariri using ttest_rel).
